# The fractionation dependence of tumor control in proton therapy for early-stage non-small cell lung cancer

**DOI:** 10.1101/2025.01.23.632803

**Authors:** Jeho Jeong, Vicki T. Taasti, Andrew Jackson, Zeno A. R. Gouw, Charles B. Simone, Philippe Lambin, Joseph O. Deasy

## Abstract

**Purpose:** The relative biological effectiveness (RBE) of tumor control for proton beam therapy (PBT) compared to photon radiotherapy (RT) is typically assumed to be independent of fractionation. To test this, we modeled published PBT outcome results for early-stage non-small cell lung cancer (NSCLC) treatments across a range of fractionation schedules.

**Materials and Methods:** All published and analyzable cohorts were included (399 patients, 413 treated lesions). Two models were used to fit the data: a previously published tumor simulation model that fits photon RT results of NSCLC across all fractionation regimes and the Fowler LQ model with a kick-off time term. The treatment effect of each cohort was referenced to the photon equivalent dose through mechanistic model simulations in a 2 Gy/weekday scenario, with radiobiological parameters determined to simultaneously best-fit all fractionation results. The tumor control RBE of each published treatment schedule, compared to the modeled photon RT effect of the same schedule, was then estimated.

**Results:** For cohorts whose treatments lasted less than three weeks (i.e., 12 fractions or less), the RBE of PBT was in the range of 1.08 to 1.11. However, for fractionated treatments stretching over four weeks or more (20-25 fractions), the relative effectiveness was much lower, with RBEs in the range of 0.82-0.89. This conclusion was unchanged using the simpler Fowler LQ + time model.

**Conclusions:** The proton RBE for hypo-fractionated schedules was 20-30% higher than for conventional schedules. The derived radiobiological parameters of PBT differ significantly from those of photon RT, indicating that PBT is influenced differentially by radiobiological mechanisms which require further investigation.

## 1. Introduction

The recent widespread introduction of proton beam therapy (PBT) continues to be used for an increasing number of cancer treatments, mainly due to the physical properties of proton beams that, in particular, include a range beyond which the radiation dose is negligible [1,2]. For example, high-dose PBT is now often used for lung cancer treatments, especially for centrally located lesions or reirradiation treatments, where the treatment efficacy of photon radiotherapy (RT) might be limited due to unavoidable overlaps of exiting beams with nearby critical organs [3–5]. However, the precise radiobiological mechanisms of PBT are underexplored and rely primarily on preclinical animal models augmented by *in-vitro* studies.

As a charged particle, protons have some fundamental differences from photons in the pattern and intensity of microscopic energy deposition, resulting in an increase in DNA local damage patterns that are more complicated than those from photon beams [6,7]. The relative biological cell kill difference of PBT can be quantified in terms of the relative biological effectiveness (RBE), which is defined as the dose of one type of radiation divided by the dose from another for the same biological effect.

In clinical practice, a nominal RBE of 1.1 is typically assumed for PBT based on *in-vitro* and *in-vivo* measurements in various conditions [8]. Although there are debates about the use of this constant RBE, and suggestions have been made to modify it as a function of depth of penetration to reflect changes in microscopic linear energy transfer [9], the constant value of 1.1 has been assumed for cooperative group clinical trials [1,10,11].

Nonetheless, there are other reasons to question a single value for the RBE of proton therapy. Compared to conventional photon therapy, proton irradiation produces, on average, DNA lesions of increased complexity and higher spatial clustering, as well as increased numbers of reactive oxygen species (ROS). These factors contribute to more highly clustered DNA repair foci and slower repair kinetics [12]. The cellular preference for homologous recombination repair (HRR) may be greater following proton irradiation [13]. Differences in the molecular processing of DNA damage may lead to differences in cell cycle arrest and cell fates, including apoptosis [14]. Alterations may also extend to immune responses [15]. Based on these differences, and perhaps others not yet recognized, we hypothesized that the clinical RBE for tumor local control may not be independent of fractionation.

To test this hypothesis, we fit the published outcome data with two models. We refitted a mechanistic tumor response model that was previously validated for early-stage non-small cell lung cancer (NSCLC) [16–18] with the clinical outcomes of PBT. This allows us to compare radiobiological parameters for PBT and also estimate the RBE for any fractionation regimes. To probe the robustness of these conclusions, we also modeled PBT outcomes using the much more straightforward and well-known LQ plus time approach popularized by Fowler.

## 2. Materials and Methods

### 2.1. Clinical outcome datasets

A literature search was performed through PubMed using the following terms: “proton & (local control OR LC) & (SBRT OR SABR OR hypofraction*) & lung”; 41 papers were found. The resulting papers and relevant references within the papers were reviewed for inclusion, among which nine papers had enough information for model simulation and outcome analysis [19–27]. The model simulation requires a detailed fractionation schedule with a distinct outcome. The reasons for excluding the other datasets include: overlapping patient cohort with currently included data, heterogeneous dose-fractionation in a combined outcome, review (meta-analysis) papers, no outcome available, and no information about fractionation schedules.

From the nine papers, twelve patient cohorts were identified with distinct local control rates reported, comprising a total of 399 patients (413 lesions). Treatment details and outcomes were collected, including fractional dose, number of fractions, fractionation schedule and overall treatment duration, and local control (LC) rate at 2 years. When the LC was reported at a different time point, the 2-y LC rate was determined from survival curves at 2 years.

The twelve cohorts were categorized into three different fractionation groups to assess if there was a fractionation dependency: 20-25 fx, 10-12 fx, or 5 fx or less. For all cohorts, the proton dose was reported in photon equivalent dose (Gray equivalents (GyE) or cobalt Gray equivalents (CGE)), assuming a generic proton RBE of 1.1. The physical dose of PBT was derived by dividing the reported equivalent dose by the RBE of 1.1. Detailed information on the included patient cohorts is summarized in Supplementary data (Table S1).

### 2.2. Tumor response model simulation and EQD2_model_ estimation

A previously developed mechanistic tumor response model was used to simulate the tumor response to PBT and correlate it with the conventional photon dose for comparison [16–18]. Briefly, the model is based on the classical radiobiological mechanisms, including the linear quadratic cell-kill model (LQ-model), hypoxia, repopulation, reoxygenation, and cell cycle variability of radiosensitivity, which are well-recognized as essential factors in tumor response to radiation. The model also incorporates an ‘energy budget’ concept that considers a local competition for chemicals (oxygen and glucose) that are necessary for cell survival and proliferation. The local chemical supply was assumed to be constant throughout radiotherapy, and this determines the number of cells in three different states: proliferating (P), intermediate (I), and hypoxic (H). Due to the different availability of nutrients and oxygen, cells in different states have different cell proliferation and death dynamics, as well as different radiosensitivity, as shown in Figure 1.

**Figure 1.**
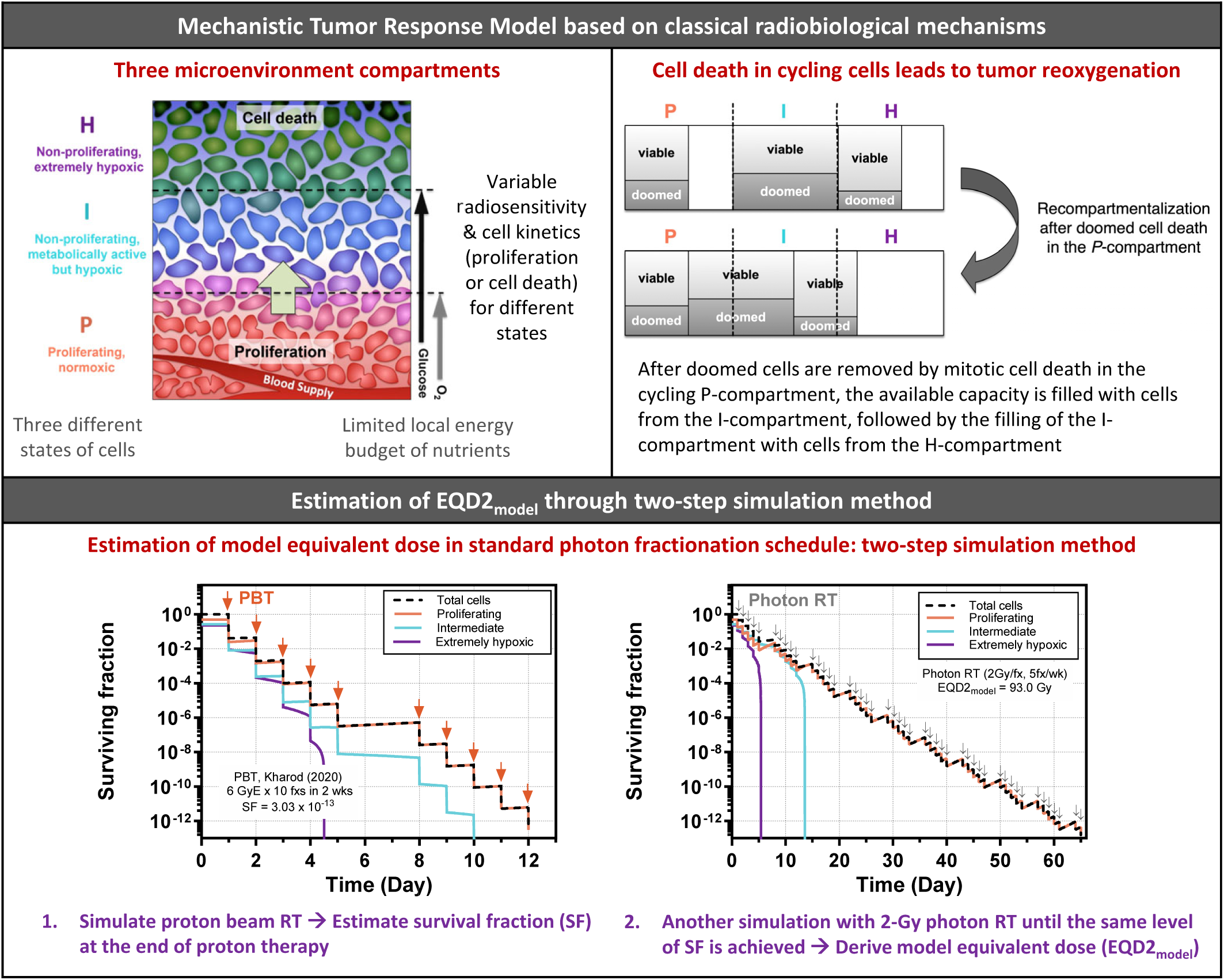
Brief description of the mechanistic tumor response model (upper) and the two-step simulation method to estimate model equivalent photon dose in 2Gy/weekday fractionation schedule (EQD2_model_) (bottom).

To compare the outcomes of different modalities (PBT vs. photon RT) and various fractionation schedules, the tumor cell-kill effect of each patient cohort was normalized based on a model-derived dose metric: the model-equivalent photon dose in conventional fractionation of 2Gy/weekday (denoted EQD2_model_). To derive the EQD2_model_ for each cohort, a two-step simulation method was used. First, a model simulation was performed for a given PBT schedule with proton-specific radiosensitivity values to find a resulting cell survival fraction (SF). Then, another simulation was run for conventional photon RT schedules (2Gy/weekday) with pre-determined photon parameter values [17], until the SF became equivalent to that of PBT, resulting in the EQD2_model_. An example of the 10-fraction PBT cohort [24] is shown in Figure 1.

Parameter values for the model simulation are summarized in Table 1. All parameter values except for the proton radiosensitivity parameters are the same as those used in Jeong *et al*. [17].

**Table 1.**
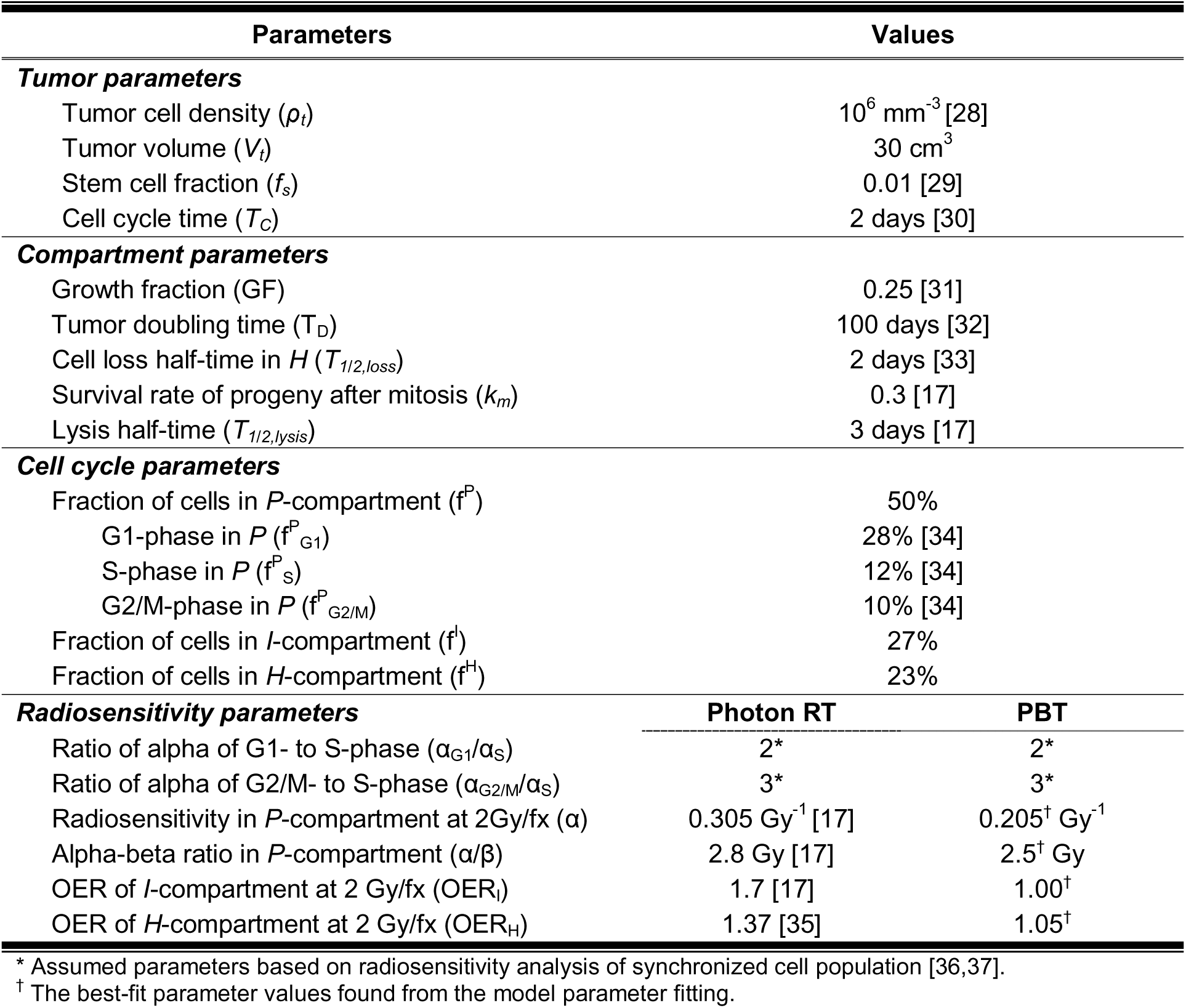
The parameters used for the model simulation for early-stage lung cancer. The numbers in brackets after values are the references.

### 2.3. Model parameter fitting

In the previous photon outcome data analysis, we found a representative dose-response relationship for early-stage lung cancer in terms of EQD2_model_ [17], which was used for reference in the current study. The tumor dose at which a tumor control probability (TCP) of 50% is expected (TD_50_) was found to be 62.1 Gy (in EQD2_model_), with the slope of the dose-response curve (γ_50_) of 1.5. The upper bound of the dose-response curve was assumed to be 95%, considering that the local control typically saturates at about 95%.

To see if the simple RBE factor of 1.1 can account for the biological difference between protons and photons, we first tried to fit the PBT outcome data using radiosensitivity parameters of photon RT (Table 1) with the reported photon equivalent dose of protons (in CGE or GyE). Then, the radiosensitivity parameters, including α value, α/β ratio, and oxygen enhancement ratio (OER) values (for the *I*-compartment, OER_I_, and the *H*-compartment, OER_H_), were optimized to find proton-specific parameter values, with which the proton outcomes fit well on the previously derived dose-response curve. Note that in the simulation with proton-specific parameters, the physical dose of protons (i.e., the photon equivalent dose in CGE or GyE divided by 1.1) was used, instead of the photon equivalent dose.

All four parameter values were concurrently optimized until best-fit parameter values were found based on the maximum log-likelihood method [38]. The 95% confidence intervals (95% CI) of the parameter values were found through the profile likelihood method [38,39].

### 2.4. Estimation of relative biological effectiveness

As the cell-kill effect of each patient cohort was normalized into a standardized dose metric (EQD2_model_), a comparison of treatment efficacy among the various PBT fractionation schedules is possible. The ratio of EQD2_model_ over physical PBT dose (EQD2_model_/D_phy_) quantifies the cell-kill efficacy of a given PBT fractionation schedule, relative to the conventional 2Gy/weekday photon RT.

The RBE is dependent on various physical and biological conditions, including fractionation schedule, LET, tissue or cell type, and biological endpoints [10,40]. Therefore, the comparison should be performed in identical conditions, other than the differences caused by the type of radiation. As the cell-kill effect of each PBT cohort was normalized in terms of EQD2_model_, a direct comparison between PBT and photon RT was possible. To derive the RBE, the equivalent photon dose, resulting in the same EQD2_model_ as PBT with the matching fractionation schedule, was found for each PBT cohort. This was possible due to the extensive validation of the model for photon RT schedules [17]. Then, the RBE was derived from the equation below:

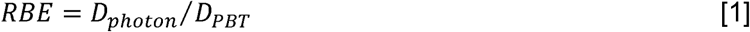

Where, *D_PBT_*is the total physical dose of PBT and *D_photon_* is the total photon dose that results in the same EQD2_model_ as PBT for the matching fractionation schedule.

The variability of the RBE value was tested by varying the fitting parameters (α value, α/β ratio, and OER values) within the 95% confidence intervals, and the range of the RBE was estimated for each PBT cohort.

### 2.5. Fowler’s LQ + Time formula

Fowler’s LQ + Time equation was also used to independently fit the outcome data and estimate the RBE for different fractionation groups. We previously showed that the LQ + time equation can precisely fit the photon outcome data, similar to the mechanistic radiobiological modeling [41]. The Fowler LQ + time equation calculates the log cell-kill effect based on the LQ model as well as the repopulation effect with a repopulation term that does not start until the kick-off time (T_k_), as shown below [42]:

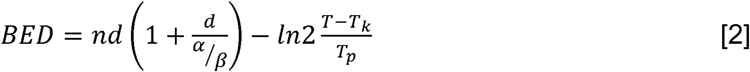

where, *n* is the number of fractions, *d* is the fractional dose in photon equivalent dose (GyE or CGE), α*/*β is the alpha-beta ratio (=10 Gy), *T* is the overall treatment time, *T_k_* is the kick-off time of fast proliferation (=28 days), α is the linear radiosensitivity parameter (=0.35 Gy^-1^), and *T_p_*is the average tumor doubling time (=3 days).

The RBE for each fractionation group was estimated by comparing the centroid of outcome data points of that group against the dose-response curve representing the photon RT outcome.

## 3. Results

To test if the generic RBE of 1.1 is adequate to describe PBT outcome data, we first applied the photon parameters with PBT dose in CGE or GyE. Figure 2 shows the comparison of outcome data fitting between photon RT vs. PBT through the model simulation (upper row). Also, the fitting through Fowler’s LQ + time formula was compared (lower row). Both comparisons indicate that PBT outcome fittings are not very successful with lower p-values of 0.77 and 0.86 compared to the photon data fitting. Although the fit for hypofractionation groups (10-12 fx and 5 fx or less) was reasonable, the treatment efficacy of the 20-25 fx group was overestimated. Optimizing the OER values in the model simulation did not improve the fit (Figure S1).

**Figure 2.**
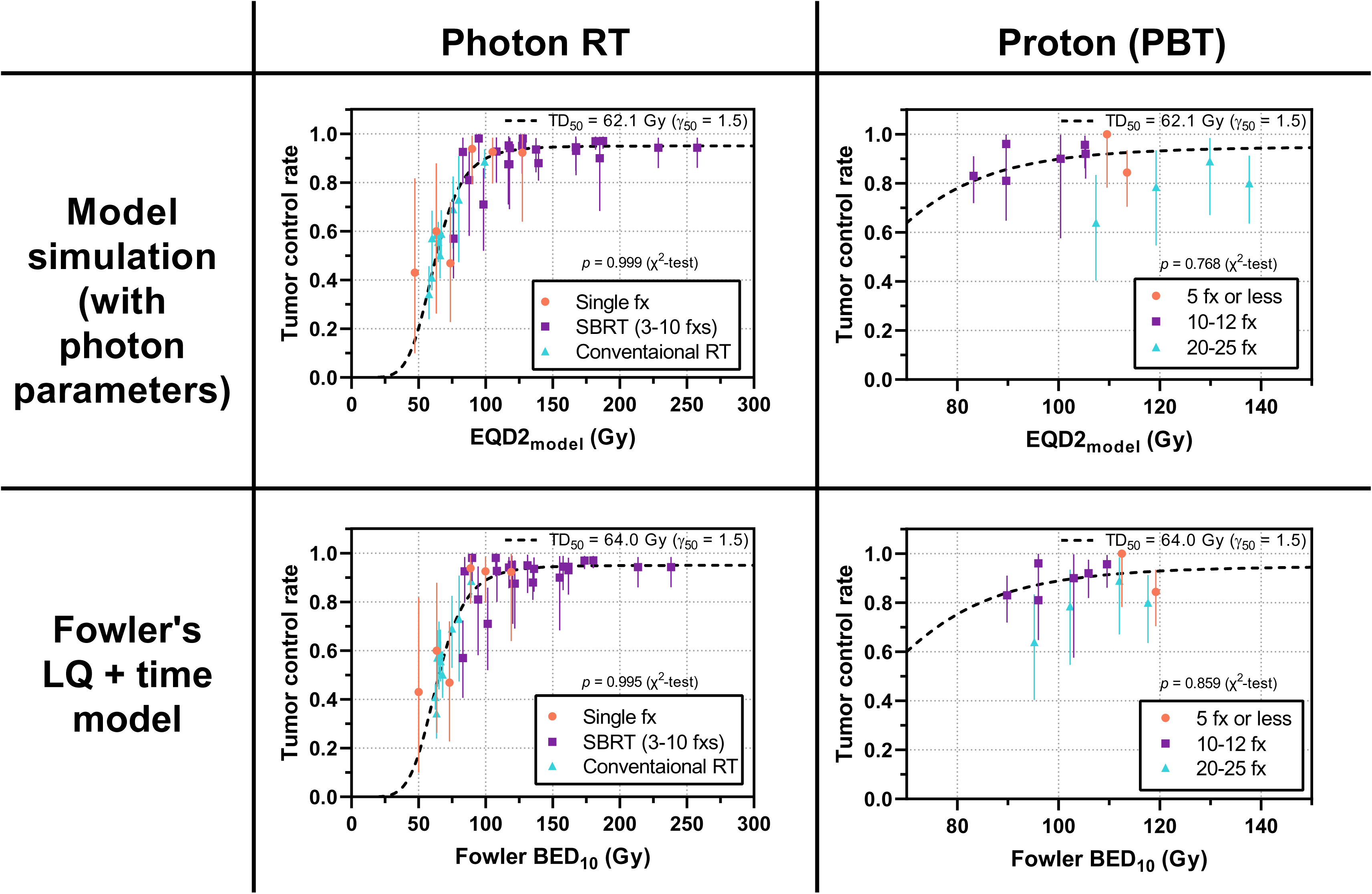
Testing the validity of generic proton RBE value of 1.1. Compared to the excellent fit of photon outcome data for both model simulation and Fowler’s LQ + time model (left column), the PBT outcome fittings are less robust with lower p-values (right column). The error bars of TCP indicate the 95% confidence interval (95% CI) based on the Clopper-Pearson method.

The radiobiological model parameters of PBT were optimized to fit the outcome data, and the best-fit parameter values and uncertainty range were found: α = 0.205 Gy^-1^ [95% CI: 0.100-0.320], α/β ratio = 2.5 Gy [95% CI: 0.9-5.6], OER_I_ = 1.00 [95% CI: 1.00-1.37], and OER_H_ = 1.05 [95% CI: 1.00-1.60]. Note that the photon equivalent dose (proton physical dose × 1.1 in CGE or GyE) was used when the photon parameter was applied in Figure 2 and Figure S1, while physical proton dose was used when the proton specific parameter values were derived in Figure 3(a).

**Figure 3.**
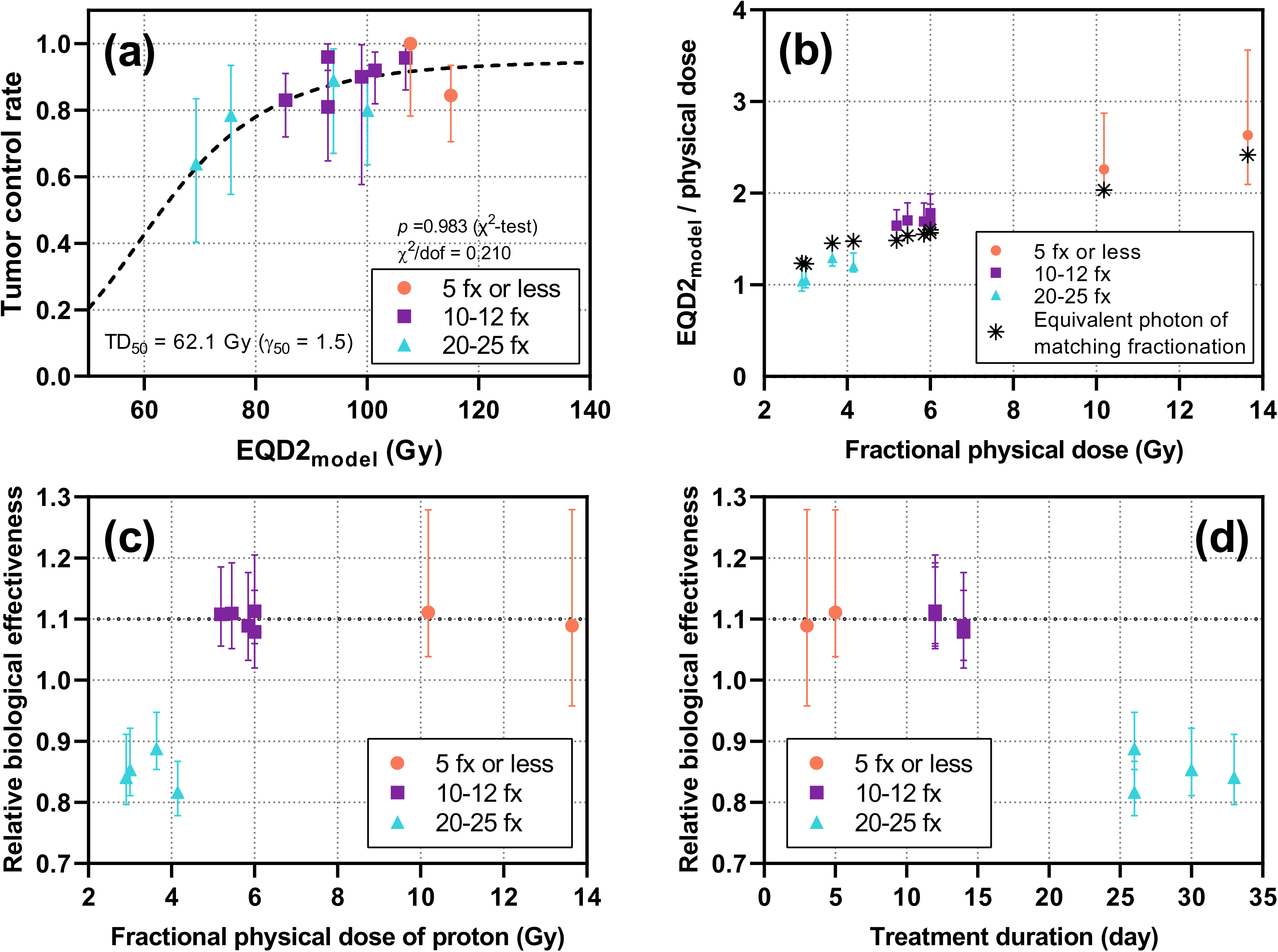
The PBT outcome data fitting, relative treatment efficacy expressed in EQD2_model_/D_phy_, and relative biological effectiveness (RBE), derived through the optimization of model parameters: (a) Fitted PBT outcome data with best-fit parameter values (α = 0. 205 Gy^-1^, α/β = 2.5 Gy, OER_I_ = 1.00 and OER_H_ = 1.05), overlaid with the previously derived photon 2 Gy/weekday dose-response curve for early-stage lung cancer [17] (TD_50_=62.1 Gy and γ_50_=1.5, dashed curve). The resulting χ^2^ test p-value was 0.983 with a low χ^2^/dof of 0.21, indicating a good fit, the error bars of TCP indicate the 95% confidence interval (95% CI) based on the Clopper-Pearson method; (b) the EQD2_model_/D_phy_ indicates the relative treatment efficacy of PBT compared to the conventional photon RT, which increases with the increase of fractional dose; (c) the RBE is approximately 1.1 for fractional dose of 5 Gy/fx or greater (10-12 fx and 5 fx or less groups), whereas it was much lower for fractional dose less than 5 Gy/fx with longer treatment schedules (20-25 fx group); (d) the RBE vs. treatment duration showing that PBT treatment schedules longer than 3 weeks have lower treatment efficacy with lower RBE.

With the best-fit parameter values, the PBT outcome data fit very well on the dose-response curve with a high p-value of 0.983 and a low χ^2^ per degree of freedom (dof) of 0.21, as shown in Figure 3(a). The 2D log-likelihood distributions around the best-fit parameter values are shown in Figure S2, from which the 95% confidence interval for each parameter was found. The relative treatment efficacy of each PBT cohort (denoted as EQD2_model_/D_phy_), which represents a relative gain in treatment efficacy of the given PBT regimen compared to conventional 2Gy/weekday photon therapy, shows increasing trend from 1.04 to 2.64 as the fractional dose increases, as shown in Figure 3(b) and Table 2. The error bars show the variation of the ratio when the parameter values were varied within 95% confidence intervals.

**Table 2.** The relative biological effectiveness (RBE) of proton beam therapy (PBT), derived from the comparison with the equivalent photon RT dose of matching fractionation schedule.

| Groups | Fractional physical proton dose (Gy) | Total physical proton dose (Gy) | Model-derived EQD2 (Gy) | $EQD2_{model}/D_{phy}$ | Equivalent photon dose of matching fractionation (Gy) | RBE [range] | Ref |
| --- | --- | --- | --- | --- | --- | --- | --- |
| 20-25 fx | 2.91 | 72.7 | 75.6 | 1.04 | 61.2 | 0.841 [0.797-0.912] | [19] |
|  | 3.00 | 66.0 | 69.3 | 1.05 | 56.4 | 0.854 [0.811-0.921] | [20] |
|  | 3.64 | 72.7 | 94.0 | 1.29 | 64.6 | 0.888 [0.854-0.948] | [21] |
|  | 4.15 | 83.0 | 100.1 | 1.21 | 67.8 | 0.817 [0.778-0.867] | [22] |
| 10-12 fx | 5.19 | 51.9 | 85.4 | 1.65 | 57.5 | 1.108 [1.056-1.185] | [23] |
|  | 5.45 | 54.5 | 93.0 | 1.70 | 60.5 | 1.109 [1.052-1.192] | [24] |
|  | 5.45 | 54.5 | 93.0 | 1.70 | 60.5 | 1.109 [1.052-1.192] | [21] |
|  | 5.85 | 70.2 | 99.0 | 1.69 | 76.5 | 1.089 [1.033-1.176] | [25] |
|  | 6.00 | 60.0 | 106.9 | 1.78 | 66.8 | 1.113 [1.060-1.205] | [26] |
|  | 6.00 | 60.0 | 101.4 | 1.69 | 64.7 | 1.079 [1.020-1.147] | [20] |
| 5 fx or less | 10.18 | 50.9 | 115.0 | 2.26 | 56.5 | 1.111 [1.039-1.279] | [25] |
|  | 13.64 | 40.9 | 107.8 | 2.64 | 44.6 | 1.089 [0.958-1.279] | [27] |

The RBE of each PBT cohort was estimated from equation 1, in which the treatment efficacy was compared with photon RT of matching fractionation schedules. In Figure 3(b), the relative treatment efficacy of the matching photon RT was also shown in an asterisk for each cohort. For cohorts with a smaller number of fractions (12 fx or less), the estimated RBE was consistently in the range of 1.08 to 1.11, which is very close to the clinically used generic RBE value of 1.1. However, the RBE was evaluated to be much lower for longer treatment schedules (20-25 fx), in the range of 0.82-0.89. For those cohorts, the treatment efficacies of photon RT in matching fractionation schedule (asterisks in Figure 3(b)) were higher than those of PBT, resulting in RBEs lower than 1. The estimated RBE values are shown in Figure 3(c-d) and Table 2.

The trend of reduced RBE for longer fractionation schedules was further supported by Fowler’s LQ + time analysis. The estimated RBE for the longer fractionation group, consisting of 20 to 25 fractions, was found to be 0.85. In contrast, the RBE for groups with 12 or fewer fractions was closer to the clinical RBE (1.12 for 10-12 fx group and 1.09 for 5 fx or fewer group).

## 4. Discussion

To test the hypothesis that the RBE for PBT vs. photon RT varies as a function of fractionation, we applied a mechanistic tumor response model to fit the PBT outcome data into the previously validated dose-response relationship of early-stage lung cancer. To test the adequacy of photon equivalent dose (CGE or GyE) based on the generic RBE of 1.1, we first applied the photon parameters to the PBT outcome data. However, the resulting fit was unsatisfactory, with an overestimated dose response for the 20-25fx group, even if the OER values were optimized (Figures 2 and S1). This implies that the use of a constant RBE factor is not adequate to explain the clinically observed PBT outcomes in this study.

The estimated RBE values in this study were dependent on the fractionation schedule. For the larger fractional doses with shorter schedules (> 5 Gy/fx in less than 12 fx), the clinically derived RBE agrees with the generic RBE currently used in the clinic. However, for smaller fractional doses with longer schedules (< 5 Gy/fx in 20-25 fx), the evaluated RBE was significantly lower than the generic 1.1 RBE, in the range of 0.82-0.89. Although the generic RBE of 1.1 is based on extensive *in-vitro* and *in-vivo* measurements in various conditions [8], most of those measurements were conducted for high doses per fraction.

The results suggest the use of generic RBE of 1.1 might need to be revisited for various fractionation schedules, especially for smaller fractional doses. This may have considerable implications for lung cancer treatments that are commonly delivered in 30 or more fractions, such as for locally advanced NSCLC or limited-stage small cell lung cancer. It may also explain, in part, why an early trial comparing PBT and photon RT over 37 fractions for lung cancer showed no significant difference in local control [43] and may influence the outcomes of the ongoing RTOG 1308 randomized trial delivering up to 35 fractions for lung cancer [44].

Despite the use of the model that worked well for photon RT, the fitted parameters of PBT cannot be accepted at face value and apparently hide some unknown effects. We could get an excellent fit to the proton data with the optimized radiosensitivity parameters, as shown in Figure 3(a). When the derived parameters for PBT were compared to those of photon (Table S2), however, it was surprisingly found that the fitted α value was lower than the photon value (0.205 Gy^-1^ vs. 0.305 Gy^-1^). In addition, the fitted OER values were significantly different from the photon values and were close to 1.0. In the model, the α and α/β ratio represent the radiosensitivity of non-hypoxic cells (in the *P*-compartment), whereas the OERs represent the relative radiosensitivity of hypoxic cells.

In both respects, the modeling results are unexpected, as they represent cells becoming more resistant to proton irradiation while discounting the impact of hypoxia. Unsurprisingly, we conclude that something is missing from our model. We briefly discuss two areas we suspect contribute to the observed difference.

Recent research emphasizes that DNA repair pathway preferences of PBT are altered compared to photon irradiation [45–47]. While non-homologous end joining (NHEJ) is an essential repair process for any radiation source, PBT is more dependent on HRR due to its more complex and condensed damage with a higher LET [12,13,46]. Therefore, late S and G2-phase cells that have access to the HRR pathway could be the most resistant and determine the response to proton therapy, while non-cycling hypoxic cells have decreased synthesis of HR proteins and reduced repair capacity of clustered DNA damage without a homologous template. However, it is difficult to see how this effect could be as large as what is observed here, and we were unable to find a model emphasizing HRR that could account for the data.

A second area of interest involves cell cycle arrest, which is not explicitly captured in the current model. It has recently been shown that the length of cell cycle arrest for photon irradiation can be a strong function of dose, with larger delays resulting from higher fraction doses [48]. It has also been found that PBT induces a more pronounced G2/M phase arrest in the cell cycle due to the greater DNA damage and higher levels of ROS compared to photon RT [14,49]. As a result, this disruption may alter the overall dynamics of the tumor, including the cell cycle time and the distribution of cells in different cell cycle phases. Theoretically, this could lead to slower overall tumor reoxygenation for PBT, which is known to be a significant factor in tumor control.

The current model does not explicitly account for those effects from HRR and cell cycle arrest. Future research could use on-treatment, cone-beam imaging to compare tumor mass loss between photon and proton therapy to detect any significant impact due to the difference in cell cycle arrest.

Despite these limitations, the fitted standardized dose index, the EQD2_model_, can normalize the treatment efficacy of any modality and fractionation schedule and enables a direct comparison of treatment efficacy among different modalities and fractionation schedules. As shown in Figure 4, the clinical outcome data from three different modalities (photon, proton, and carbon) in various fractionation schedules can be fit well into a single dose-response curve for early-stage lung cancer, in terms of EQD2_model_.

**Figure 4.**
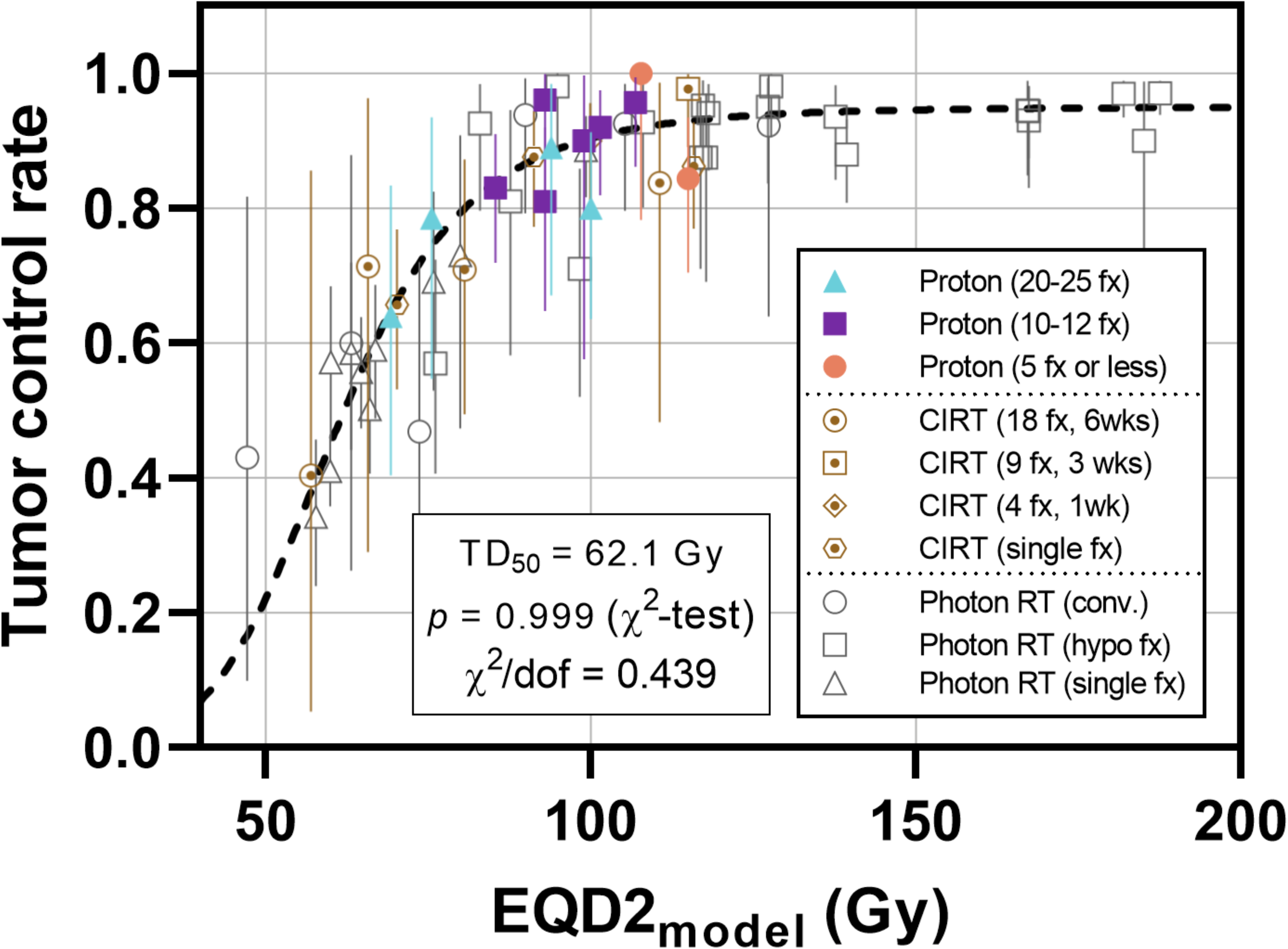
The combined early-stage lung cancer outcome data from PBT, carbon ion radiation therapy (CIRT), and photon RT, overlaid with the previously derived dose-response curve (TD_50_=62.1 Gy and γ_50_=1.5). All data fit well in a single-dose response curve with a very high χ^2^ test p-value. Photon RT and CIRT results are from Jeong et al. [17,18]. The error bars of TCP indicate the 95% confidence interval (95% CI) based on the Clopper-Pearson method.

Some important limitations apply to this research, besides those discussed above, which include: (1) the amount of PBT clinical data, and the variety regarding fractionation, is limited, (2) it was not possible to fold LET differences into the analysis, (3) we only had adequate data for analysis for the single site of early-stage NSCLC, and (4) the model is clearly a simplified representation of the complicated reality of tumor response and some important details in modeling are apparently missing. Despite these caveats, the observation that PBT results depart from photon RT results is robustly supported using either the tumor simulation model or the empirical LQ + time model popularized by Fowler (see Figure 2).

## 5. Conclusions

For early-stage non-small cell lung cancer, the relative biological effect of proton beam therapy regarding tumor control is about 1.1 for hypo-fractionated therapy, but drops to 0.8-0.9 for fractionation regimes delivered over more than about three weeks. Validating this result and understanding the radiobiological source of the difference will require further research.

## Acknowledgments

This study was supported by the MSK Cancer Center Support grant (P30 CA008748), the Simons Foundation, a Breast Cancer Research Foundation grant (MATH-23-001), and NIH grant U54CA274291.

