## Supplementary material for "The fractionation dependence of tumor control in proton therapy for early-stage non-small cell lung cancer": Table S1, Table S2, Figure S1, and Figure S2

### Supplementary data

**Table S1. Detailed patient cohort data included in this study (additional columns on the next page).**

| **Fraction Group** | **#** | **Ref** | **Institution** | **Treatment duration** | **# pt** | **# lesion** | **Age (yr)** | **Gender (M/F)** | **T-stage**  **(T1/T2)** | **Tumor location** | **Beam arrangement** | **proton energy (MeV)** | **Motion management** |
| --- | --- | --- | --- | --- | --- | --- | --- | --- | --- | --- | --- | --- | --- |
| 20-25 fx | 1 | Ono, 2017 [19] | Southern Tohoku | 01/2009-02/2015 | 20 | 20 | med: 75 (63-90) | 17/3 | 4/11 (T3/4=4/1) | Central | 1-3 portals;  passive scattering | 150/210 | CT gated at exhalation/ gating for end-expiration |
|  | 2 | Kanemoto, 2014 [20] #1 | U of Tsukuba #1 | 02/1997-09/2011 | 21 | 21 | med 75 (51-86) | 65/15* | 59/21 | Central | 2-3 beams | 155-250 | gating for expiratory phase |
|  | 3 | Iwata, 2010 [21] #1 | HIBMC at Tatsuno #1 | 04/2003-04/2007 | 20 | 20 | med 75 (48-87) | 13/7 | 6/14 | NR | 1-4 portals | 150 | CT gated at exhalation/ gating for end-expiration |
|  | 4 | Nihei, 2006 [22] | NCC in Chiba | 12/1999-10/2003 | 37 | 37 | med 75 (63-87) | 30/7 | 17/20 | NR | 2-4 portals | 150-190 | gating for expiratory phase |
| 10-12 fx | 5 | Bush, 2004 [23] | Loma Linda | NR | 68 | 68 | mean: 72 [52-87] | 30/38 | 29/39 | NR | 3-4 beams | NR | fluoroscopy/margin added to account for respiratory motion |
|  | 6 | Kharod, 2020 [24] | U of Florida | 10/2009-8/2018 | 23 | 24 | med 74 (58-88) | 11/12 | 11/13 | Central | 3-4 beams; double scattering | NR | 4DCT/iGTV |
|  | 7 | Iwata, 2010 [21] #2 | HIBMC at Tatsuno #2 | 04/2003-04/2007 | 37 | 37 | med 78 (57-87) | 30/7 | 21/16 | NR | 1-4 portals | 150 | CT gated at exhalation/ gating for end-expiration |
|  | 8 | Lee, 2016 [25] #1 | PTC in Korea #1 | 09/2007-12/2013 | 11 | 11 | med 75 (47-89) | 43/12* | 39/16 | Central | 2-5 (median3); coplanar or non-coplanar beams | 230 | 4DCT/gate if tumor motion is more than 1cm (29 pt (53%)) |
|  | 9 | Hatayama, 2016 [26] | Southern Tohoku | 01/2009-09/2014 | 50 | 52 | med 72.5 (54-87) | 35/15 | 44/8 | Peripheral | 2-3 portals | NR | CT gated at exhalation/ gating for end-expiration |
|  | 10 | Kanemoto, 2014 [20] #2 | U of Tsukuba #2 | 02/1997-09/2011 | 53 | 59 | med 75 (51-86)* | 65/15* | 59/21 | Peripheral | 2-3 beams | 155-250 | gating for expiratory phase |
| 5 fx or less | 11 | Lee, 2016 [25] #2 | PTC in Korea #2 | 09/2007-12/2013 | 44 | 44 | med 75 (47-89)* | 43/12* | 39/16 | Peripheral | 2-5 (median3); coplanar or non-coplanar beams | 230 | 4DCT/gate if tumor motion is more than 1cm (29 pt (53%)) |
|  | 12 | Westover, 2012 [27] | MGH | 07/2008-09/2010 | 15 | 20 | med 78 (62-89) | 3/12 | 18/2 | 18 peri /  2 cent | 2-3 beams (coplanar); passive scattered | NR | 4DCT/mid-ventilation approach |
| NR: not reported.  *the values are for total patient cohort reported in the paper, not separated by different fractionation groups | | | | | | | | | | | | | |

**Table S1. Detailed patient cohort data included in this study (continued).**

| **Fraction Group** | **#** | **Total dose (GyE)** | **Dose/fx (GyE)** | **# of fraction** | **Physical total dose (Gy)** | **2y-LC (%)** | **EQD2_model_** | **Equivalent photon dose**  **of matching fractionation (Gy)** | **Proton EQD2_model_ /D_phy_** | **EQD2_model_ /D_phy_ of the matching photon** | **RBE** |
| --- | --- | --- | --- | --- | --- | --- | --- | --- | --- | --- | --- |
| 20-25 fx | 1 | 80 | 3.2 | 25 | 72.7 | 78.5 | 75.6 | 61.2 | 1.04 | 1.24 | 0.841 |
|  | 2 | 72.6 | 3.3 | 22 | 66.0 | 63.9 | 69.3 | 56.4 | 1.05 | 1.23 | 0.854 |
|  | 3 | 80 | 4.0 | 20 | 72.7 | 89.0 | 94.0 | 64.6 | 1.29 | 1.46 | 0.888 |
|  | 4 | 70/80/ 88/94 | 3.5/4.0/ 4.4/4.7 | 20 | 63.6/72.7/ 80.0/85.5 | 80.0 | 100.1^†^ | 67.8 | 1.21 | 1.48 | 0.817 |
| 10-12 fx | 5 | 51/60 | 5.1/6 | 10 | 46.4/54.5 | 83.0 | 85.4 | 57.5 | 1.64 | 1.48 | 1.108 |
|  | 6 | 60 | 6.0 | 10 | 54.5 | 96.0 | 93.0 | 60.5 | 1.70 | 1.54 | 1.109 |
|  | 7 | 60 | 6.0 | 10 | 54.5 | 81.0 | 93.0 | 60.5 | 1.70 | 1.54 | 1.109 |
|  | 8 | 60/72 | 6.0 | 10/12 | 54.5/65.5 | 90.0 | 99.0^†^ | 63.7 | 1.69 | 1.55 | 1.089 |
|  | 9 | 66 | 6.6 | 10 | 60.0 | 95.7 | 106.9 | 66.8 | 1.78 | 1.60 | 1.113 |
|  | 10 | 66 | 6.6/5.5 | 10/12 | 60.0 | 92.0 | 101.4^†^ | 64.7 | 1.69 | 1.57 | 1.079 |
| 5 fx or less | 11 | 50/60 | 10.0/12.0 | 5 | 45.5/54.5 | 84.4 | 115.0^†^ | 56.5 | 2.26 | 2.03 | 1.111 |
|  | 12 | 45 | 15.0 | 3 | 40.9 | 100.0 | 107.8 | 44.6 | 2.63 | 2.42 | 1.089 |
| 2y-LC: 2-year local control rate; EQD2_model_: model equivalent dose in photon 2Gy/weekday fractionation; RBE: relative biological effectiveness.  ^†^population averaged EQD2_model_ value for various fractionation schedules included in the cohort | | | | | | | | | | | |

**Table S2. Comparison of the best-fit radiobiological parameter values with the 95% confidence interval for three different types of radiation (proton, photon, and carbon ion).**

| **Parameters** | **Best-fit value [95% CI]** | | |
| --- | --- | --- | --- |
|  | **Proton** | **Photon [17]** | **Carbon [18]** |
| α value | 0.205 [0.100-0.320] | 0.305 [0.12-0.365] | 1.12 [0.97-1.26] |
| α/β ratio | 2.5 [0.9-5.6] | 2.8 [0.4-4.4] | 23.9 [8.9-38.9] |
| OER_I_ | 1.00 [1.00-1.37] | 1.70 [1.55-2.25] | 1.08 [1.00-1.41] |
| OER_H_ | 1.05 [1.00-1.60] | 1.37* | 1.01 [1.00-1.44] |

* Not optimized.

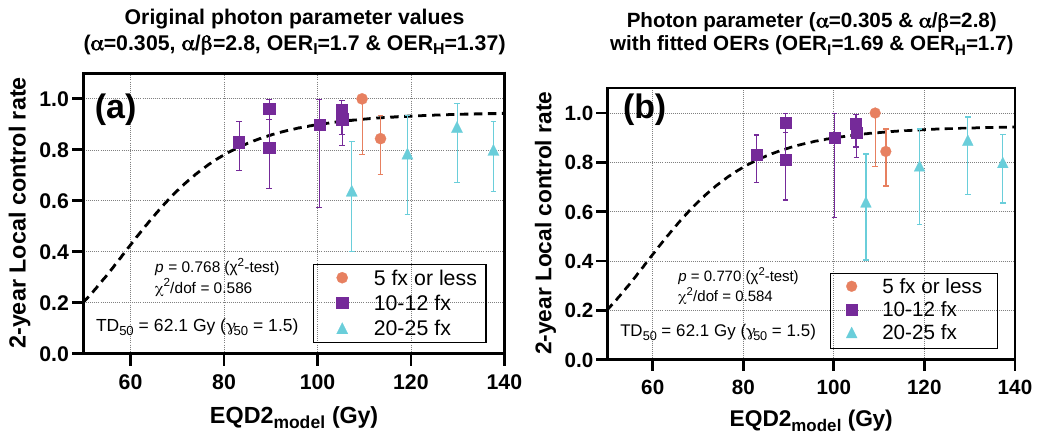

**Figure S1. Initial model fit of PBT outcome data with parameter values derived from photon RT, overlaid with the previously derived photon 2 Gy/weekday dose–response curve for early-stage lung cancer [17] (TD_50_ = 62.1 Gy and γ_50_ = 1.5, dashed curve): (a) original photon parameter values (α = 0.305 Gy^-1^, α/β ratio = 2.8 Gy, OER_I_ = 1.70, and OER_H_ = 1.37); (b) OER values were optimized for best-fit, keeping α value and α/β ratio. In either case, the dose-response of the 20-25fx group was overestimated, resulting in an unsatisfactory fit with p-values around 0.77. The error bars of TCP indicate the 95% confidence interval (95% CI) based on the Clopper-Pearson method.**

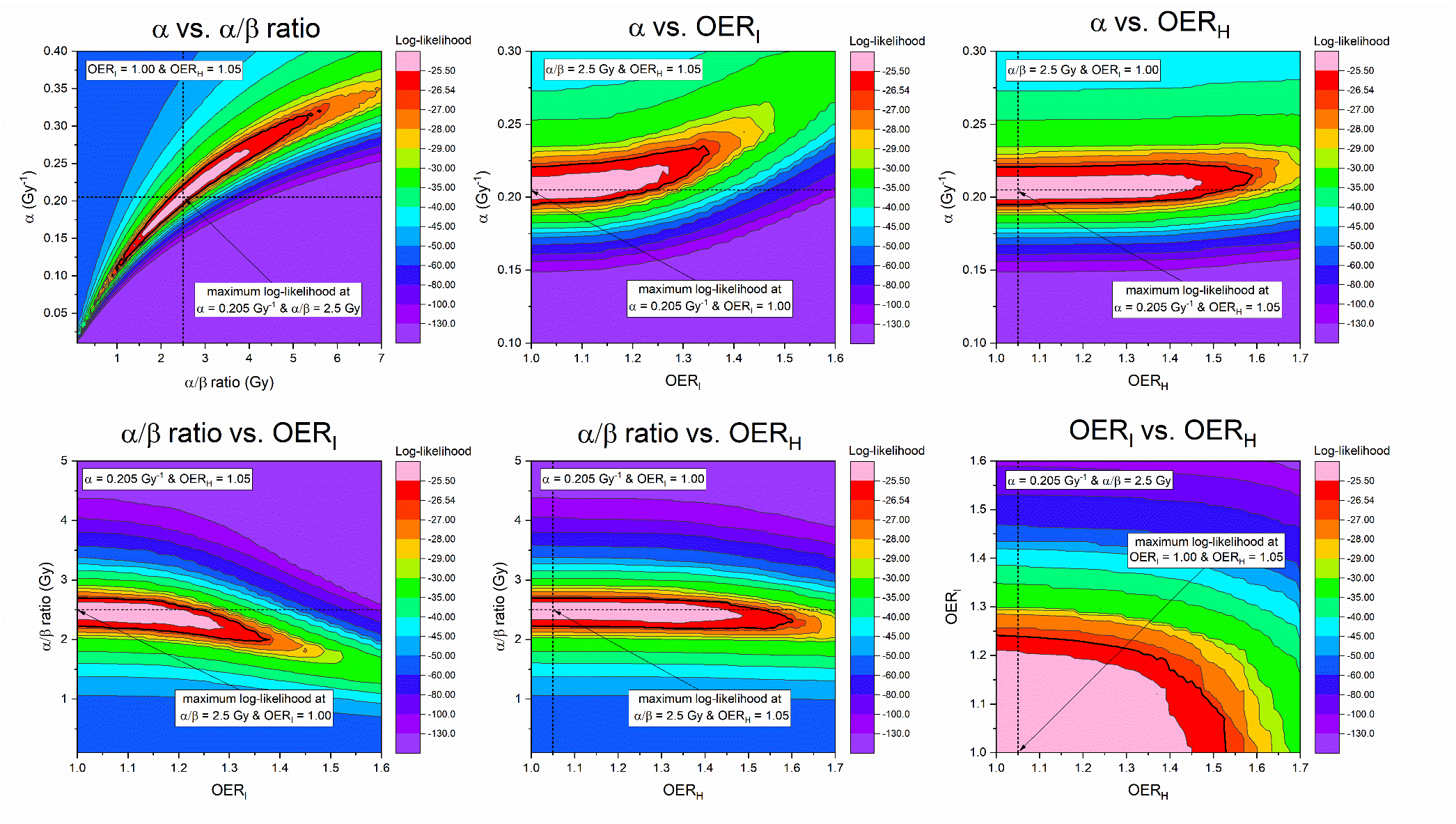

**Figure S2. 2D cross-sectional views of log-likelihood distribution around the best-fit parameter values. The thick black contour in each plot indicates the 2D 95% confidence intervals (95% CI) found through the profile likelihood method.**
